# Movement-related drivers of exposure to West Nile virus by American robins (*Turdus migratorius*)

**DOI:** 10.1101/2023.10.27.562968

**Authors:** Alex E. Jahn, Kyle Koller, Lynn B. Martin, Tara M. Smiley, Taylor B. Verrett, Ellen D. Ketterson, Emily J. Williams, Daniel J. Becker

**Affiliations:** Environmental Resilience Institute, Indiana University, Bloomington, IN, USA; Department of Biology, Indiana University, Bloomington, IN, USA; Global Health Infectious Disease Research Center, College of Public Health, University of South Florida, Tampa, FL, USA; Department of Ecology and Evolution, Stony Brook University, Stony Brook, NY, US; School of Biological Sciences, University of Oklahoma, Norman, OK, USA; Department of Biology, Georgetown University, 37th and O Streets NW, Washington DC, USA Washington, DC 20057, USA

**Keywords:** bird, disease ecology, dispersal, migration, Midwest

## Abstract

The ecological processes that determine how individual animals become hosts to zoonotic pathogens is a topic of rapidly growing interest. However, how such exposure is mediated by context (e.g., season, location), host behavior (e.g., migration distance) and host demographics is generally poorly understood. We evaluated seasonal exposure to West Nile Virus of American robins sampled in Indiana and compared our results to those of previous studies. Because robins that breed in Indiana are partial migrants (i.e., only a portion of the population migrates), we evaluated their probability of exposure to WNV as a function of whether they migrated or not and of their movement distance. We also tagged a subset of breeding robins with tracking devices to evaluate their potential to disperse the virus between Indiana and other regions of the continent. We found that robins that breed in Indiana are exposed to WNV at a higher rate than that detected in previous studies, but found no correlation between robin exposure and whether a robin migrated or not, nor with migration distance, season, sex, and breeding latitude (for robins overwintering in Indiana). Our tracking data indicate that robins that breed in Indiana migrate several hundred miles to overwinter in the southeastern US. The mean duration of their return to Indiana in spring is 10.3 days, which is less than the maximum infectious period found for robins in previous studies, suggesting that they have the capacity to move WNV long distances in spring. However, we still know little about the physiological capacity of robins to migrate while being infectious, which could inhibit the dispersal of the virus through robin migration. Future research on the physiological, ecological and behavioral factors mediating the exposure of birds to WNV will lend insight into the role that robins and other birds play in the transmission ecology of the virus.

## Introduction

Rapid environmental changes are modifying ecosystems globally, including the relationship between wildlife and their pathogens (Hassell et al. 2017; Murray et al. 2019). Emerging infectious diseases can severely negatively impact wild bird populations (Robinson et al. 2010; Dadam et al. 2019), and wild birds can also serve as competent hosts for common zoonotic pathogens, including but not limited to *Borrelia spp.* (Becker and Han 2020; Richter et al. 2000), Rickettsia spp. (Lommano et al. 2014; Becker et al. 2022), Anaplasma spp. (Keesing et al. 2012), *Babesia spp.* (Hersh et al. 2012), and West Nile virus (WNV) (Komar et al. 2003; Kilpatrick et al. 2006).

Nevertheless, we largely lack a conceptual framework of the ecological and evolutionary processes that determine which birds are more likely to host and disperse pathogens, including the suite of physiological and life history characteristics that mediate these processes. In large part, this is due to generally better sampling during the spring breeding season relative to the rest of the year. Birds may invest differentially into body condition and immune function throughout the year as a function of different movement strategies (e.g., that require changes in energy expenditure) in different seasons (Hegemann et al. 2012). As a result, they may have different probabilities of exposure and of becoming infected across seasons (Altizer et al. 2006).

WNV appeared in North America in 1999 (Nash et al. 2001), causing mortality in humans, horses, and birds (LaDeau et al. 2007, Gould et al. 2017) and which has spread across much of the continent through the migration of birds (reviewed by Komar et al. 2001, Reisen 2013). The magnitude of the viremia in at least some migratory birds does not appear to change significantly during migration (Owen et al. 2006), lending further support to the idea that birds are the main dispersers of this virus. However, we know almost nothing about the sex-specific risk of infection to WNV in birds, which is key because immune function is often dependent upon the trade-offs faced by different sexes (Klein 2000, reviewed by Martin et al. 2008). Nor do we understand how variation in susceptibility to WNV may change throughout the year, which may also vary by sex (Altizer et al. 2006) and result in major epizootics (e.g., in American crows, *Corvus brachyrhynchos*; Townsend et al. 2023).

Susceptibility is also likely to be seasonally- and regionally-specific; for example, a population breeding at lower latitudes may be exposed to WNV-carrying mosquitoes during more months of the year than a population breeding at higher latitudes, where the onset of winter is earlier (reviewed by Hoover and Barker 2016). Additionally, migratory birds may have a higher probability of exposure to pathogens as a function of the distance they move (López et al. 2008). Although migratory birds likely play an important role in the long-distance (i.e., both intra- and inter-continental) dispersal of WNV (e.g., López et al. 2008, Llopis et al. 2015, Mancuso et al. 2022), we still understand little about the mechanisms that promote or constrain the virus’ dispersal.

American Robins are considered a “superspreader” of WNV because they are regularly fed upon by the mosquito vectors of WNV (Apperson et al. 2004, Kilpatrick et al. 2006, Hamer et al. 2009, Simpson et al. 2012), develop viremia (VanDalen et al. 2013) and are competent for WNV (Komar et al. 2003). Movements of robins have also been linked to probability of their exposure to the virus, since their late summer/early fall dispersal away from breeding sites to communal roosting sites makes them less available to mosquito vectors of WNV (Kilpatrick et al. 2006, Janousek et al. 2014). However, the relationship between robin movements and occurrence of WNV in mosquito vectors appears to be related to regional- and year-effects (Benson et al. 2012). Because year-long sampling of robins for WNV is lacking, our ability to evaluate season-specific predictors of pathogen positivity in robins is limited.

Urban adapters or exploiters, such as the American Robin (Turdus migratorius), can benefit from suburban landscapes, where they forage on invertebrates and fruits that are common in lawn and ornamental plant ecosystems (McKinney 2002). Such species often have greater reproductive success, survival, and abundance in suburban habitats as compared to rural habitats (Rodewald and Bakermans 2006; Schneider et al. 2015; Morneau et al. 1995; Evans et al. 2015). All else being equal, such as how occupancy in urban habitats affects exposure and susceptibility to pathogens (Murray et al. 2019; Becker et al. 2015), the greater overlap between bird hosts, arthropod vectors, and humans in these habitats should facilitate cross-species transmission.

Our goal was to evaluate seasonal variation in exposure of American Robins (hereafter, “robins”) to WNV in a Midwestern urban area by comparing WNV seroprevalence levels in robins sampled in Bloomington, IN between seasons and sexes. Because robins are partial migrants in IN and other Midwestern states of the USA (i.e., some robins migrate while others remain on their breeding site year-round; Vanderhoff et al. 2020), they may exhibit a wide range of seasonality in behavior and physiology that could affect their exposure to WNV, which are still poorly understood in migratory birds.

To address these gaps, we sampled robins throughout two years in Bloomington, IN and used serology to evaluate seasonal exposure to WNV and whether probability of exposure is related to whether a robin is a migrant or a resident (i.e., non-migratory). We tested the hypothesis that, for robins sampled in Indiana in winter, there would be a negative relationship between WNV seropositivity and breeding latitude, since WNV is found up to ∼55° N in North America (Hoover and Barker 2016), whereas robins in North America can breed as far as 70° N (Vanderhoff et al. 2020). We also evaluated the potential for robins sampled during the breeding season to disperse WNV to and from the Midwest by tracking a subset of migratory robins. Finally, we used formal meta-analysis to compare the seroprevalence of robins we sampled to that of robins sampled elsewhere in North America. With these results, we offer new insights and avenues for research on the ability of birds to host and disperse WNV under rapidly changing environmental conditions.

## Methods

### Bird capture and sampling

Robins were sampled on the campus of Indiana University and nearby neighborhoods in Bloomington, IN. Robins begin arriving in Bloomington on spring migration in March, then breed from April to July, although the majority of nests are active April to May (AEJ, pers. obs). Robins then undergo feather molt from August to October, and migrate south beginning in October, although some remain throughout winter (AEJ, unpub. data). Our aim was to sample robins to assay exposure to WNV during four seasons: early spring, breeding, fall, and winter. We therefore sampled robins monthly from January-May 2020, Sep-Dec 2020, and January, March and April 2021; i.e., representing one fall season (September-December 2020), two winter seasons (January-February 2020, January 2021), two early spring seasons (March 2020, 2021), and two breeding seasons (April-May 2020, April 2021). All capture sites (n = 16) were located within a 5 km radius and were composed of mowed lawns with scattered trees, intersected by buildings and sidewalks.

We captured robins using up to 5 polyester mist nets (3x18 m, 60 mm mesh; Avinet Research Supplies) from pre-dawn until late morning, under appropriate weather conditions (i.e., without rain, excessive heat or high wind). Upon capturing a robin, we banded it using a USGS numbered band, then aged and sexed it based on presence of juvenile plumage, a brood patch or cloacal protuberance (Pyle 1997). We measured unflattened wing chord and body mass (Ralph 1993) derived a measure of body condition through the residuals of a linear mixed model regressing wing chord on body mass (capture site as a random effect) in R using the lme4 package (Labocha et al. 2012). We collected a blood sample by puncturing the brachial vein with a sterile insulin needle and extracting blood with a 70 μL capillary tube (Owen 2011). Blood was stored in an empty microcentrifuge tube in a cooler for up to five hours before being spun in a centrifuge for five minutes at 3000 rpm. Plasma was drawn off using a micropipette and transferred to a microcentrifuge tube, followed by long-term storage at -20 C until serology.

Before release, we outfitted a subset of robins (n=21) with GPS tags (model PP-10 or PP- 50, Lotek Wireless, Inc.) during spring 2020 and outfitted an additional 32 robins in spring 2021. We attached tags to robins using a leg-loop harness made of 0.7 mm diameter jewelry elastic cord (StretchMagic brand). The combined weight of the PP-10 tag and harness was 1.2 g and that of the PP-50 and the harness was 2.3 g, representing less than 3% of the mass of the birds on which they were deployed (Geen et al. 2019). In no case was there evidence of the harness or tag causing injury to the birds on which they were deployed. As an additional measure of robin movement, in 2020 we collected the first secondary feather from captured robins, which was stored in a paper envelope for subsequent stable isotope analysis. Because robins molt their flight feathers on their breeding grounds before initiating fall migration (Vanderhoff et al. 2020), the stable isotope signature in those feathers is a reliable estimate of where they breed.

All captures, sampling, and tagging were conducted in accordance with authorization from the Indiana University Institutional Animal Care and Use Committee (protocol #18-28) and necessary permits (Federal Bird Banding Permit #20261, Indiana Department of Natural Resources Scientific Purposes License #20-528).

### WNV serology

Our serological assay protocol was modified from the Florida Department of Health IgM Antibody Capture Enzyme-Linked Immunosorbent Assay Protocol. Details are provided as Supplementary Information.

### Movement data analysis

GPS tags were programmed to collect data every three days (model PP-50) or every five days (model PP-10). Data were downloaded from each recovered tag using the ‘SWIFT fixes’ option in the PinPoint Host application (Lotek Wireless, Inc.), and in no case was a clock drift of >40 minutes detected.

### Feather stable isotope assays

We collected the first secondary feather (i.e., the most distal secondary feather from the right wing) for stable hydrogen isotope analyses at the Indiana University Stable Isotope Research Facility. Whole feathers were cleaned of external oils and contaminants using a 2:1 chloroflorm: methanol solution. We controlled for the effect of exchangeable hydrogen by forcing isotopic equilibration with a water vapor of known isotopic composition in a flow-through chamber system at 115°C (Schimmelmann, 1991; Sauer et al., 2009). Equilibrated feather samples were analyzed in replicate using a thermal conversion elemental analyzer coupled with a ThermoFinnigan Delta Plus XP isotope ratio mass spectrometer. δ^2^H values are reported in standard per mil notation (‰) relative to Vienna Standard Mean Ocean Water using two reference materials: USGS77 (polyethylene powder) and hexatriacontane 2 (C_36_ n-alkane 2). Analytical precision was ± 2.2‰ for δ^2^H values. Reported values reflect the isotopic composition of the non-exchangeable hydrogen per sample (Schimmelmann, 1991; Sauer et al., 2009).

Based on δ^2^H_feather_ values, we performed geographic assignments of individual birds during the breeding months using the assignR package in R (Bowen et al., 2014; Ma et al., 2020). The growing season precipitation isoscape from Bowen et al. (2005) was rescaled to a feather isoscape using known values from members of a ground-foraging guild, including *Turdus migratorius, Hylocichla mustelina, Catharus fuscescens, Catharus guttatus, and Catharus ustulatus* (Hobson et al., 2012). The recalibration model found an intercept value of -23.4‰ and explained 70% of the isotopic variation in resident individuals. Geographic assignment was estimated based on posterior probability density maps. From these maps, we retained the cells within the top 10% highest posterior probability using the qtlRaster function to generate ‘threshold maps’. Using the raster package (Hijmans, 2023), we extracted median, maximum, and minimum estimated latitude per individual as well as median latitude from the centroid of the estimated breeding range.

### Serological data analysis

We first used a generalized additive model (GAM) fit using the mgcv package in R (Wood 2017) to assess overall temporal dynamics in WNV seroprevalence, using a binomial response of seropositive and seronegative individuals aggregated to each sampling month over the two-year period, thin plate splines with smoothing penalty and restricted maximum likelihood (REML). We next used generalized additive mixed models (GAMMs) on individual seropositivity to partition annual and seasonal effects, including a smooth term of ordinal date (using a cyclic cubic spline), fixed effects of year and bird sex, and a random effect of site. We fit two separate models with an additional smooth term (thin plate spline with smoothing penalty) of either body condition or fat score, as these two covariates were strongly correlated (ρ = 0.39, p < 0.001). To assess potential lag effects between bird physiological state and WNV seroprevalence, we derived monthly mean body condition and fat score and then derived the cross-correlation between each monthly measure time series and monthly seroprevalence (Shumway and Stoffer 2000). Strong, positive correlations around the zero lag would support synchrony between mean physiological state of birds and WNV seroprevalence (i.e. in phase).

We next conducted two smaller analyses to test hypotheses about the interactions between WNV exposure and robin migratory strategies. First, for our subset of birds sampled following fall migration and prior to spring migration (October through December, January through March) and with corresponding geographic assignments for breeding grounds from δ^2^H_feather_ values (n = 50), we fit a GAMM predicting WNV seropositivity as a function of smoothed estimated breeding latitude (thin plate spline with smoothing penalty) using REML, with site again as a random effect. Lastly, for our subset of recaptured tagged birds with GPS data, we used a t-test with unequal variance to assess differences in migratory distance (log_10_- transformed) as a function of seropositivity at the time of tag deployment.

### Meta-analysis of seroprevalence

Lastly, we used a meta-analysis to compare our annual WNV seroprevalence estimates of robins in Indiana to robins from elsewhere in North America. We searched Web of Science, PubMed, and Google Scholar with the following string: (Turdus migratorius OR “American robin” OR “robin” OR “AMRO”) AND (“West Nile virus” OR WNV) AND (serol* OR serop* OR antibod* OR expos* OR ELISA). Using a systematic protocol (Figure S1) (Moher et al. 2009), we only included studies of WNV seroprevalence in wild robins. We excluded seroprevalence estimates based on xenodiagnosis (i.e., seroprevalence from mosquito blood meals traced to specific host species (Hannon et al. 2019; Komar et al. 2018), given that mosquito host selection is non-random and multiple vectors could feed on the same bird (Hamer et al. 2009; Chaves et al. 2010). We identified 12 studies for inclusion, from which we recorded the sampling state, month(s) and year(s), sample type (sera or blood spots), serological test (ELISA or plaque reduction neutralization test; PRNT), the serological cutoff for positivity, and total birds sampled and seropositive. Each record (n = 17) was an estimate of WNV seroprevalence in robins over space and time, with one-third of studies contributing multiple years or locations of data..

Using the metafor package, we calculated the Freeman-Tukey double arcsine transformed proportion of WNV-seropositive birds and sampling variances (Viechtbauer 2010). Meta-analytic models included weighting by sampling variance alongside an observation-level random effect nested in a study-level random effect to account for within- and between-study variance in seropositivity. To first assess heterogeneity in seroprevalence, we fit an intercept-only model with restricted maximum likelihood; estimates of the variance components were used to derive I^2^, the contribution of true heterogeneity to total variance in seroprevalence (Senior et al. 2016). We then fit four univariate models with the same random effects to test (i) whether seroprevalence in robins in Indiana differs from that in other states and whether variation in seroprevalence could alternatively be explained by (ii) different years of sampling, (iii) sample type, or (iv) serological test. Univariate models were compared given the small sample size. To compare the predictive power of these study-level moderators, we derived a pseudo-*R*^2^ measure as the proportional reduction in the summed variance components compared with an intercept-only model (López-López et al. 2014).

## Results

### Movement patterns

In spring 2021, we recovered GPS archival tags from three robins, and in spring 2022 we recovered tags from 11 robins, none of which appeared to have experienced negative effects from the tag (e.g., no lesions on the skin). Two of the robins from which we recovered tags in 2021 were seropositive for WNV upon deployment of the tag in 2020 as well as when the tag was recovered, while one bird was seronegative in both years. Three of the robins from which we recovered tags in 2022 were seropositive for WNV in both years, five were seronegative in both years, and three were unsampled for WNV antibodies. We found no significant difference between males and females in the rate, distance, and duration of spring or fall migration nor in date of departure and arrival in either season. Thus, we combined data from both sexes for further analyses.

The mean date of initiation of fall migration from Bloomington was 12 Nov (4.7 SE) and robins spent a mean 25 days (7.9 SE) on fall migration. Spring migration lasted a mean of 10.3 days (4.1 SE) and the mean arrival date back in Bloomington was 13 March (5.2 SE).

### WNV serology

Between January 2020 and April 2021, we detected seropositivity for WNV in 34% (95% CI: 0.27 to 0.41) of 179 American robins sampled in Bloomington, IN (101 in 2020 and 78 in 2021). Our initial GAM of overall temporal patterns in WNV seropositivity suggested a significant decline during our study period (χ^2^_0.84,6_ = 4.65, p = 0.02; Figure 1). However, our individual-level GAMMs detected no difference in WNV seropositivity by season (ordinal date), year, sex, or physiological state (Table 1). Cross-correlation analyses of mean monthly body condition or fat score and WNV seroprevalence likewise detected generally weak temporal associations (Figure 2), although high body condition predicted more negative seroprevalence 3–4 months later, and fat score was typically synchronized with seroprevalence. Lastly, for our subset of wintering robins with stable isotope data, we detected no association between breeding latitude and WNV serological status (χ^2^_0,6_ = 0, p = 0.44; Figure 2).

**Figure 1.**
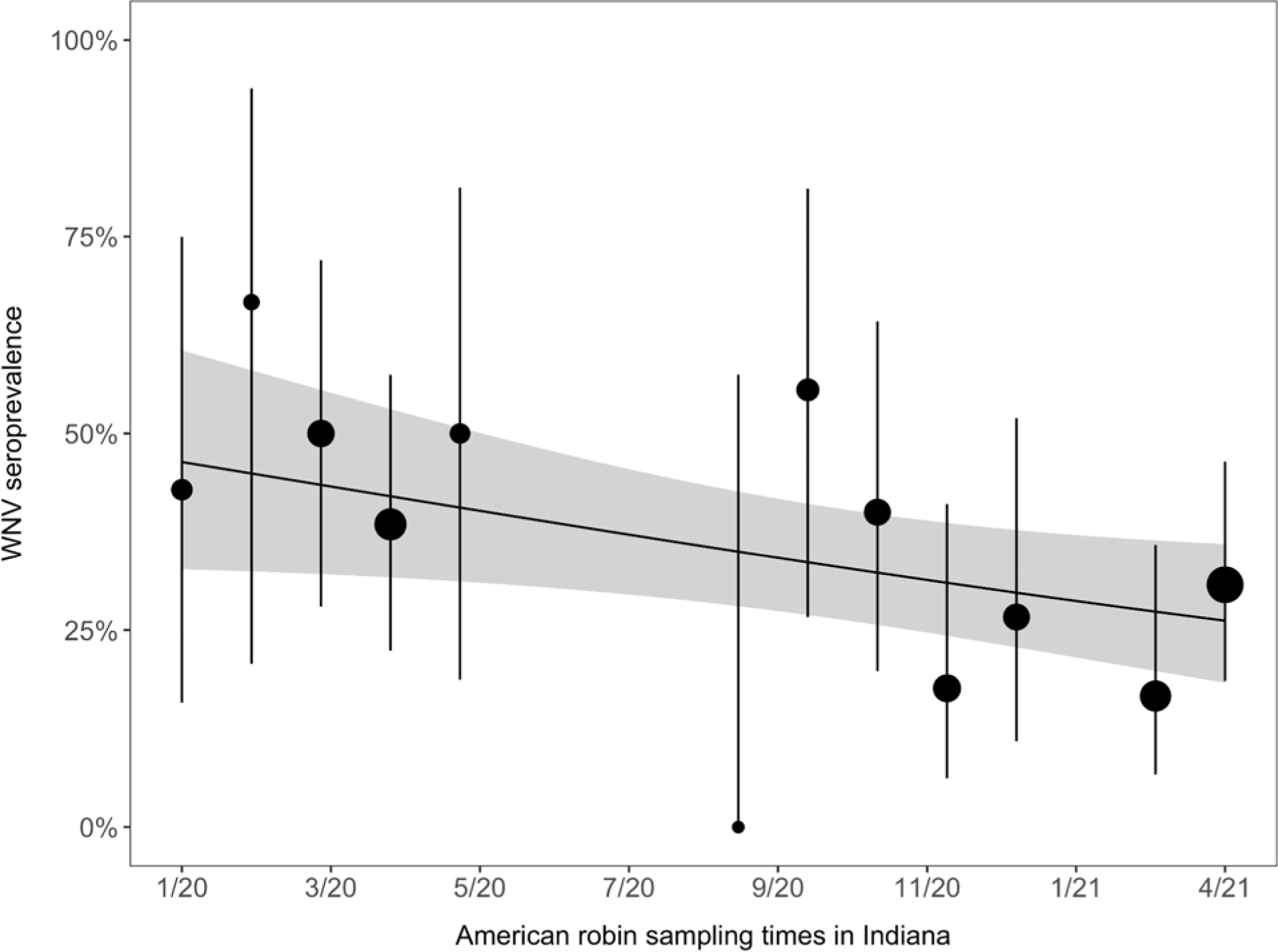
Temporal patterns in WNV seroprevalence in American robins sampled from Bloomington, Indiana. Points show seroprevalence and 95% confidence intervals (Wilson’s interval), scaled by the number of individual birds tested. The thick curve and band display the mean temporal trend and 95% confidence interval from our initial GAM.

**Figure 2.**
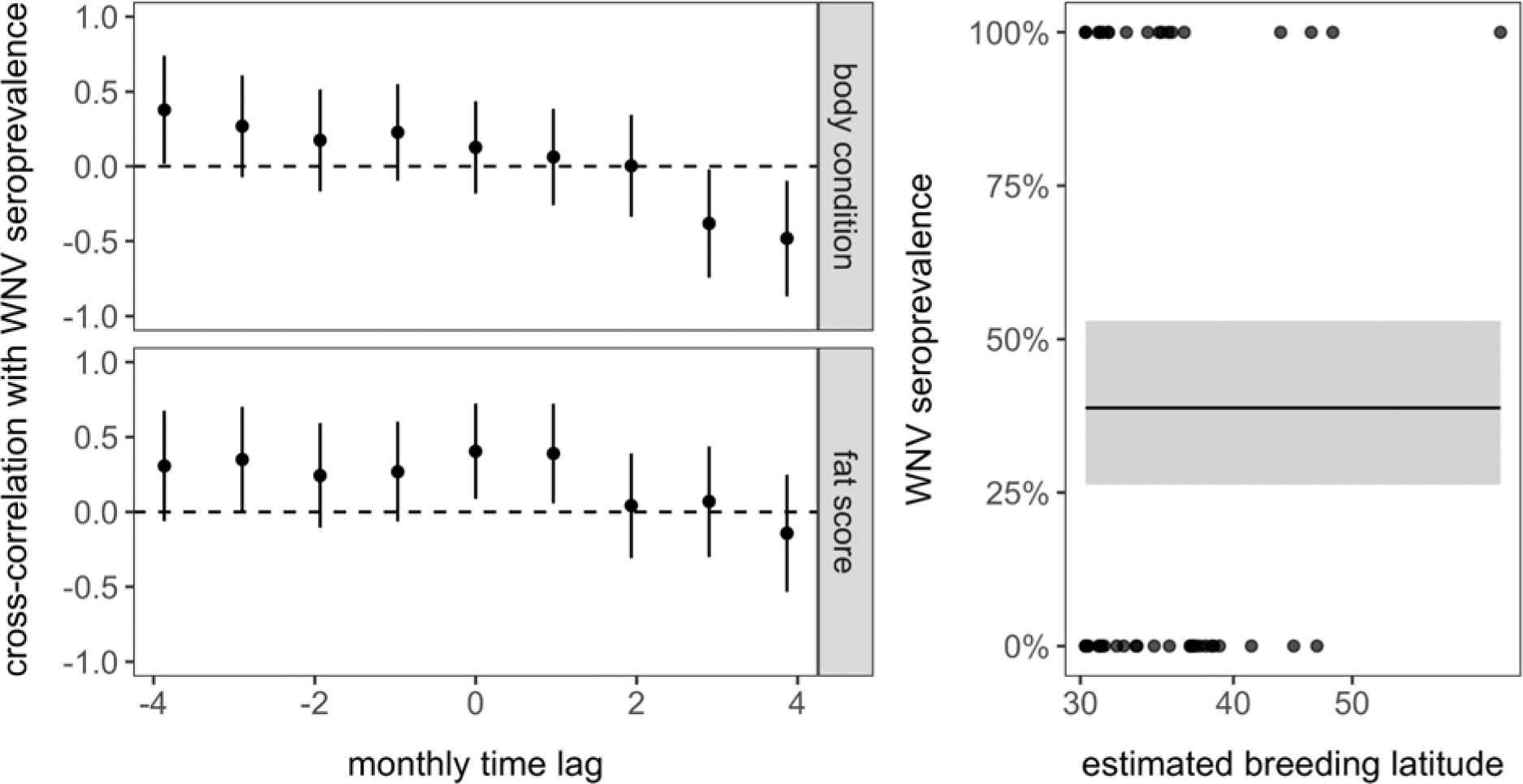
Cross-correlation coefficients and 95% confidence intervals for the lagged association between mean bird physiological state and WNV seroprevalence (A). Modeled relationship between estimated breeding latitude and WNV seropositivity (B); points show WNV seropositivity, the thick curve and band display the GAM trend and 95% confidence interval.

**Table 1.**
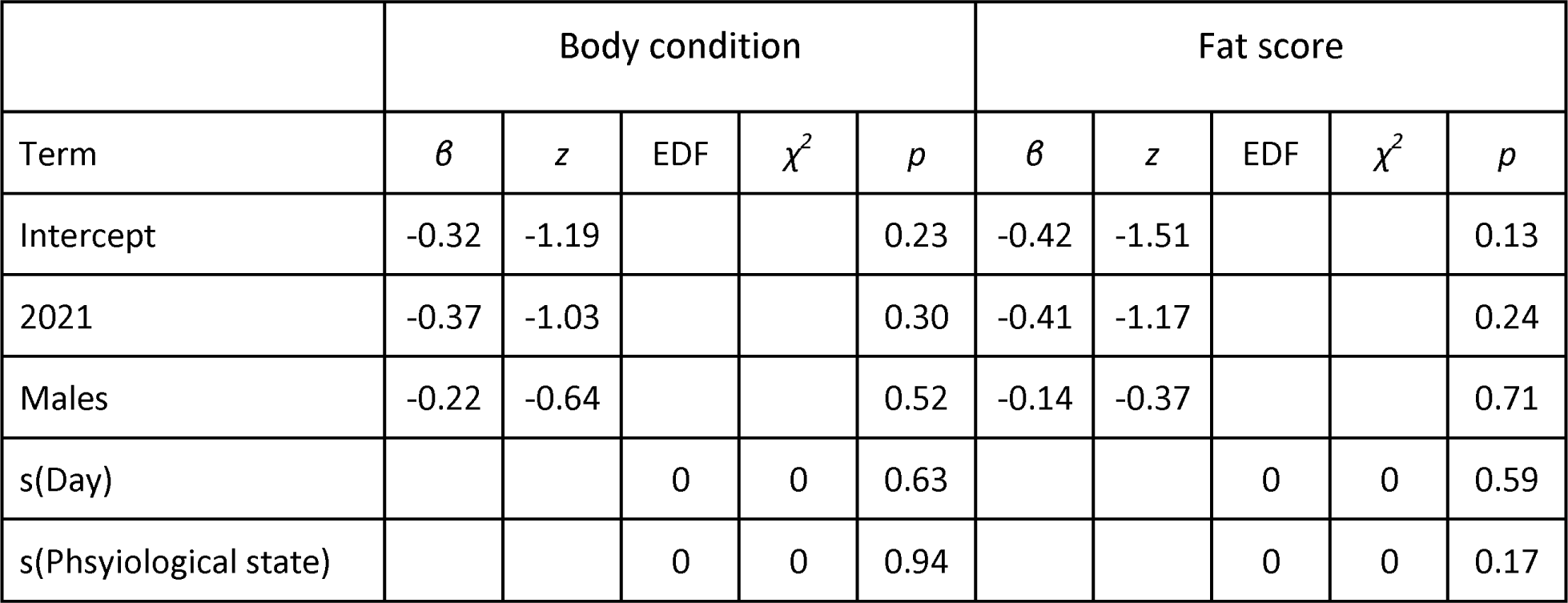
Estimated fixed effects and smoothed terms from WNV seropositivity GAMMs (site as a random effect) for models including body condition or fat score. The intercept includes birds sampled in 2020 and females.

Migraton distance (log-10 scale) did not differ between seronegatve (x̄ = 2.42, n = 6) and seropositve birds (x̄ = 2.14, n = 5; t = 0.51, df = 7.13, p = 0.63), suggesting that recent exposure to WNV (or the cost of producing WNV antibodies) has limited impact on the distance robins move (Figure 3).

**Figure 3.**
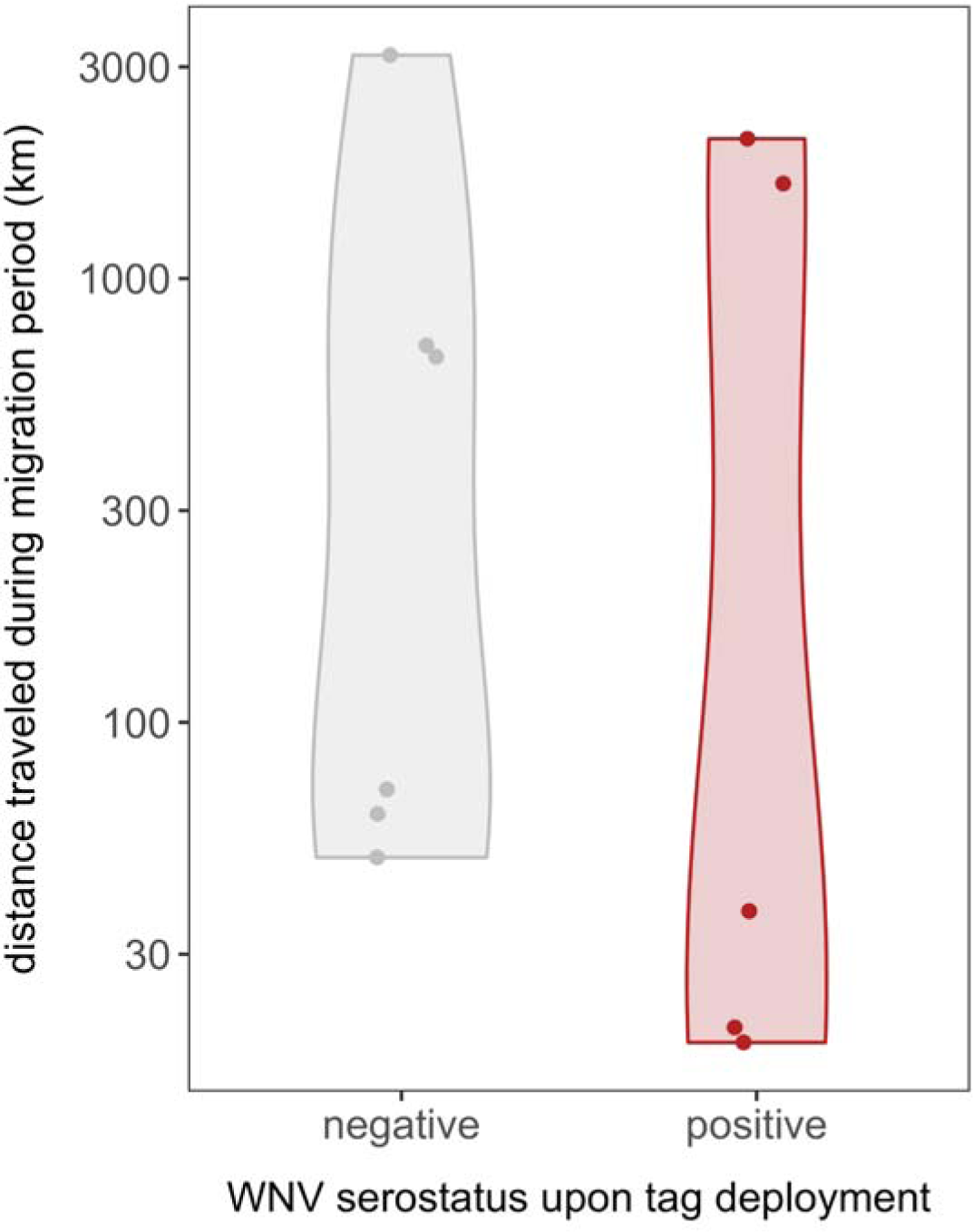
Violin plots show the distribution of migration distances as a function of WNV serological status at the time of GPS tag deployment. Raw data are overlaid and jittered.

### Meta-analysis of seroprevalence

Our systematic review of WNV serology in American robins identified 17 other seroprevalence estimates from robins sampled in New York [45–48], New Jersey [46], Louisiana [49], Arkansas [50], Georgia [51], Illinois [52–54], and Colorado [55]; one additional estimate encompassed multiple states across the Atlantic and Mississippi flyways [56]. Most estimates pooled multiple capture sites across broad spatial scales (71%), with five estimates reporting localized data. Similarly, almost all estimates spanned multiple months or did not report months of sampling (88%), with only two estimates reporting month-specific seroprevalence. Prior studies reported seroprevalence between 1999 (i.e., emergence in North America) and 2007. Our two annual seroprevalence estmates from Indiana resulted in a total of 19 records and 1444 individuals; sample sizes per estmate ranged from 1–453 birds (x̄=76, SE=24). Of these 19 estimates, only three were derived from blood spots, whereas the remainder used sera. 12 estimates used ELISAs, whereas seven estimates used PRNTs. Seropositivity cutoffs (i.e., the ability to block antibody binding or to reduce plaques) ranged from 30–45% for ELISAs and 80–90% for PRNTs; the coefficient of variation was three-times greater for ELISAs (19.17%) than for PRNTs (5.60%).

Intercept-only meta-analysis model identified substantial heterogeneity in WNV seroprevalence (Q_18_=130.11, I^2^=90.25%, p<0.001; Figure 4) and a weighted mean seroprevalence across studies of 7% (95% CI=1–17%; z=6.6, p<0.001). Seroprevalence estimates did not vary by sample type (Q_1_=1.31, β_sera_=0.18, p=0.25, *R*^2^=0) nor by diagnostic assay (Q_1_=0, β_PRNT_=-0.001, p=1, *R*^2^=0). Instead, seroprevalence increased across years (Q_1_=9.88, β=0.017, p=0.001, *R*^2^=0.64) and varied especially with geography (Q_7_=25.35, p=0.001, *R*^2^=0.71).

**Figure 4.**
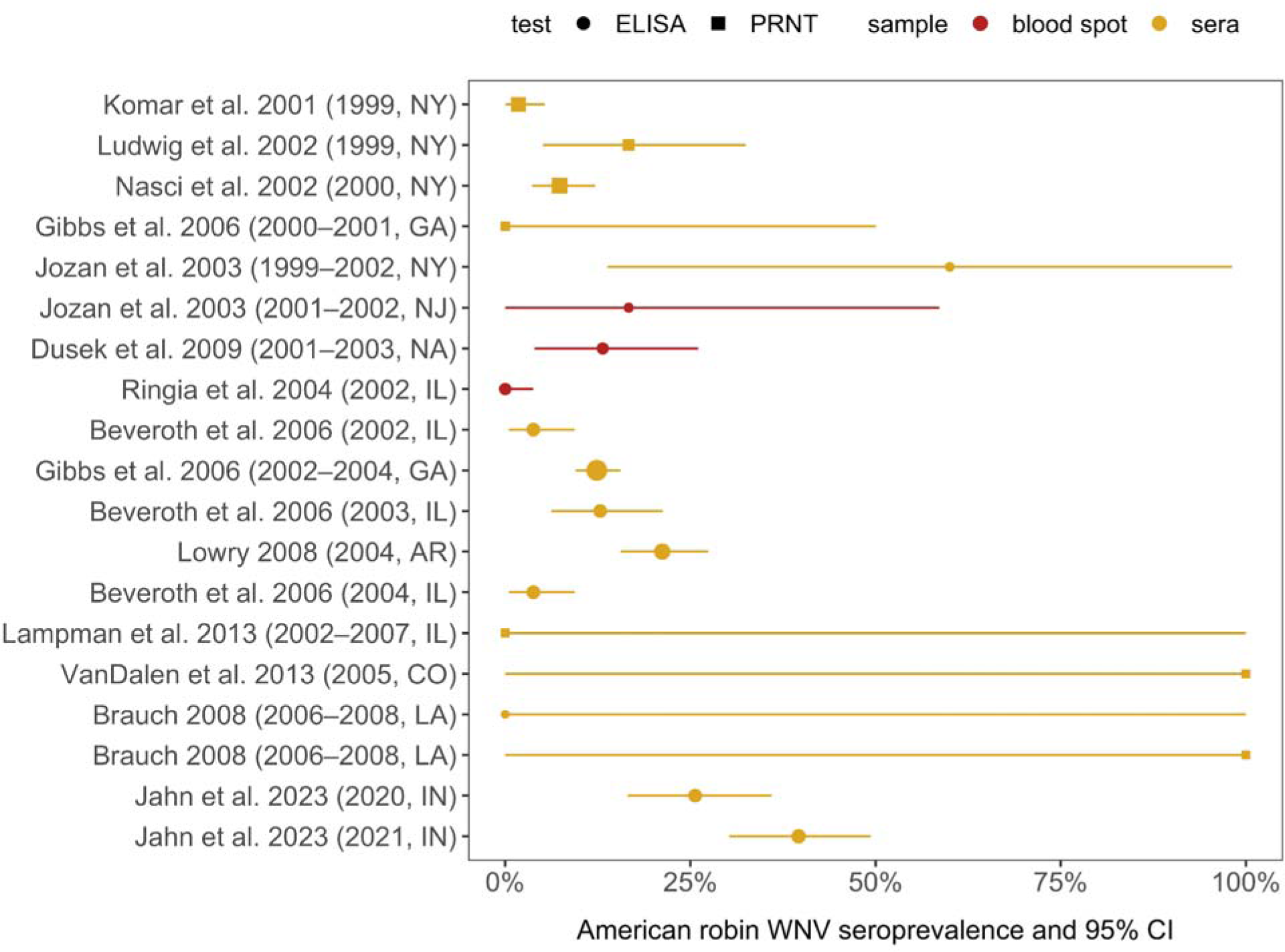
WNV seroprevalence (stratified by location and year) of American robins from systematically screened literature and associated 95% confidence intervals, including both annual estimates from our study here. Data are ordered by year, colored by sample type, shaped by serological assay, and scaled by the number of individual birds tested.

## Discussion

Our results indicate that robins in Indiana are exposed to WNV at a higher rate than that detected in previous studies in other states. Given that most previous studies were conducted over a decade ago, we cannot currently evaluate whether our results are a product of more recent higher exposure rates by robins to the virus generally/across states or if robins sampled in Indiana are an outlier. We found no correlation between robin exposure and migrant-resident status, migration distance, season, breeding latitude or sex, suggesting that the probability of exposure to WNV in the robins we sampled is a product of stochastic processes (e.g., year-specific effects) or due processes we did not evaluate (e.g., immune function). We found that a portion of robins that breed in Indiana migrate several hundred km to overwinter primarily in the southeastern US, and that exposure to WNV did not impact their ability to move, suggesting that they have the potential to disperse the virus through migration.

The duration of robin spring migration was shorter than during fall, and is similar to what has been found in studies of other migratory songbird species (e.g., McKinnon et al. 2013). Experimental studies have shown that robins develop moderately high WNV viremia and typically remain infectious for 4-12 days (Komar et al. 2003, Owen et al. 2021), which is similar to the average duration of robin spring migration that we detected (10.3 days), suggesting that the duration of infection is sufficient for the pathogen to be dispersed from robin wintering sites into the Midwest in spring. Nevertheless, spring migration may represent a dead end for WNV, since robins may arrive on a northern breeding site prior to the emergence of mosquitos at their destination. Indeed, previous studies in North America support the idea that dispersal of WNV northwards by birds is not common (reviewed by Reisen and Wheeler 2019), but recent evidence from Europe suggests northern dispersal of the virus there is plausible (García- Carrasco et al. 2021).

Even if northward dispersal of the virus is currently not viable, as climate change proceeds, mosquitos may emerge earlier at northern latitudes than they do at present (Hoover and Barker 2016). If robins respond to earlier spring emergence by advancing the date of their spring migration timing, as has been found to occur in other songbirds (e.g., Jonzén et al. 2006), and if they are able to acquire and then disperse WNV northwards during spring migration, the virus may be able to infect mosquitoes that feed on the robins when they arrive at northern breeding sites. Given that year-round transmission of WNV exists in birds along the US coastal region of the Gulf of Mexico (Dusek et al. 2009), which is the region that robins we tracked from Indiana overwinter, this scenario is plausible. Nevertheless, migration may inhibit the ability of infected individuals to spread the virus due to migratory culling, in which migrants experience pathogen-induced mortality during migration (Bradley and Altizer 2005), migratory recovery, in which birds recover from infection during migration (Shaw and Binning 2016), or migratory stalling, which is a positive relationship between parasite burden and a reduction in movement (Peacock et al. 2020). Whether these mechanisms operate in robins is as yet unknown, but given that robins are partially migratory in the Midwest, the constraints on these processes likely depend on the relative risk to WNV infection by migrants vs residents, as has been suggested for partially migratory animals generally (Shaw et al. 2023).

The potential for robins to disperse WNV also depends on the degree to which their immune function is suppressed during migration. For example, a congener of robins in Europe, the Redwing (Turdus iliacus), has been shown to suppress immune function during migration, leading to reactivation of bacterial infection (Gylfe et al. 2000). In contrast, Swainson’s thrushes (Catharus ustulatus), which are closely related to robins, exhibit migratory activity while infected with WNV and do not suppress their immune function during migration (Owen et al. 2006). Recent research suggests that food limitation can induce robins to have higher viral titers and to be infectious for a longer period (Owen et al 2021), suggesting that the physiological and ecological context is key when considering the risk of infection by robins to WNV. Research on various aspects of the transmission ecology of WNV, including the constraints on hosts, vectors and on the pathogen itself will permit a foothold on the relationship between climate and WNV transmission dynamics, which are generally poorly understood (Kramer et al. 2019).

Future sampling of robins during migration and in different conditions would provide deeper insights into the drivers and constraints of their ability to disperse WNV, since evidence exists that the types of habitat and abiotic conditions that birds use play a role in mediating the exposure of birds to pathogens, especially during migration (reviewed by Daversa et al. 2017, Shaw et al. 2023). For example, urban habitats can restructure infection risk for avian hosts (Martin and Boruta 2013), altering the composition of avian communities, which typically show an overall reduced species richness in favor of few generalists (especially omnivores and ground-foragers) that can capitalize on anthropogenic food and nesting resources (Faeth et al. 2005; Marzluff 2001). Urban birds may also exhibit a suppressed stress response (Partecke et al. 2006), although the consequences of that in terms of risk of infection throughout their annual cycle are not well understood. Given that the emergence of several avian pathogens, such as Trichomonas gallinae and WNV, have occurred in urban habitats (Lawson et al. 2012; Marra et al. 2004), this question is relevant beyond WNV.

Finally, in the context of understanding the role that pathogens play in driving bird migration strategies, sampling across conspecifics with different migration strategies (e.g., that migrate earlier vs. later or that migrate long vs. short distances) would offer novel insights into the tradeoffs faced by both migratory host and pathogen. For example, the relationship between infection intensity and migration distance can be non-linear (Shaw et al. 2023). Full annual research is therefore needed to better grasp what are likely to be a complex suite of tradeoffs between host and pathogen, mediated by the selection pressures operating on each. Comparing closely related taxa with different behavioral (e.g., migration vs residency) and ecological (e.g., habitat use) characteristics promises novel insights into how migratory animals and pathogens interact in different contexts (Shaw et al. 2023). Given the speed at which ecosystems across the planet are changing as a result of human activities, understanding how such processes are likely to impact both wildlife and humans in the near future is imperative.

## Acknowledgements

We thank the many people who provided invaluable time and effort in the lab and field, and the landowners who allowed us to work on their properties. This study was funded by the Prepared for Environmental Change Grand Challenge Initiative at Indiana University. Sampling was approved by the Indiana University IACUC (06-242, 18-028), Federal Bird Banding Permit 20261, and Indiana Scientific Purposes License #20-528.

## Supplementary Information

**Figure S1.**
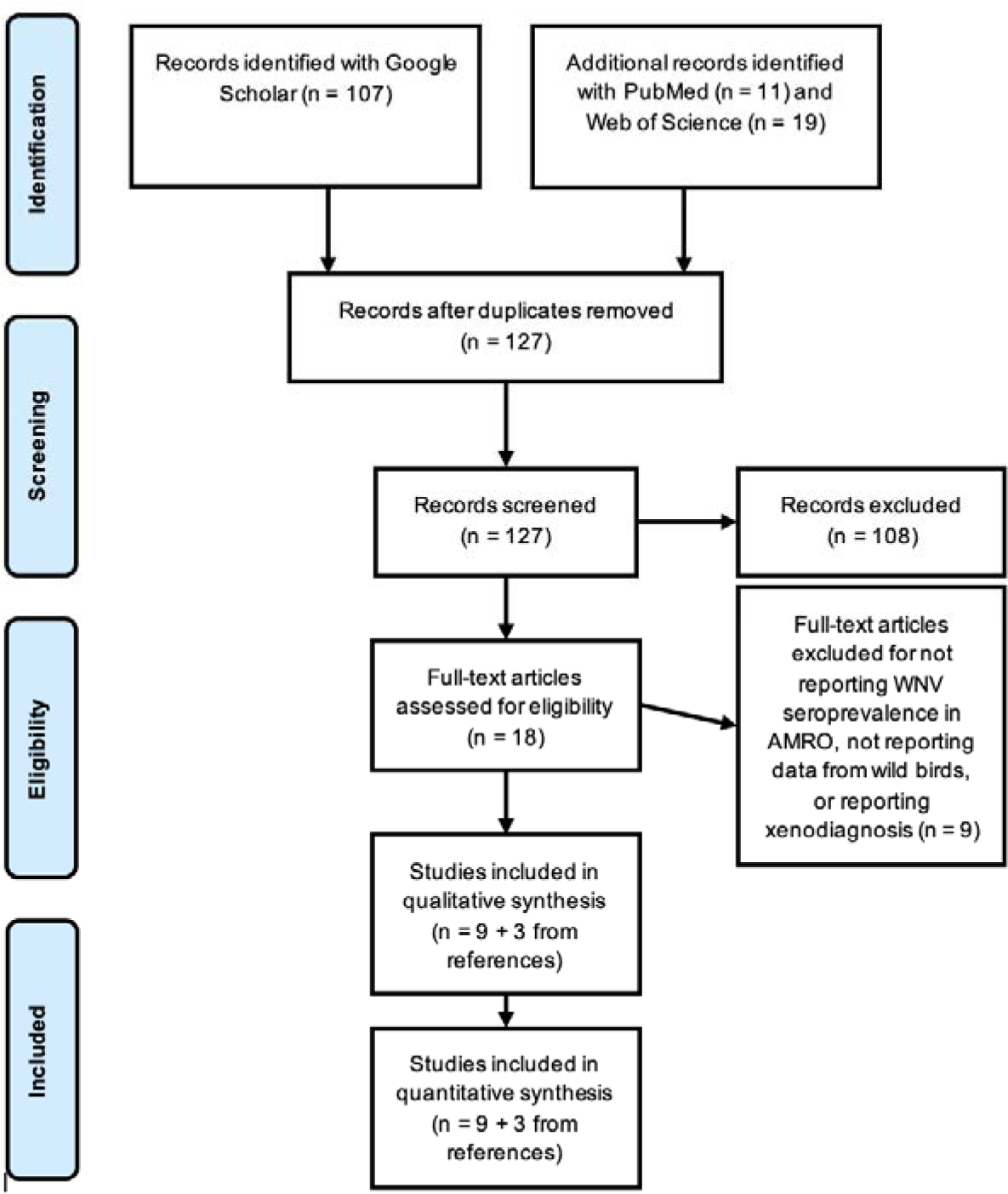
PRIMSA diagram documenting the data collection and inclusion process for WNV seroprevalence in American robins. We ran all systematic searches in July 2021 and supplemented results by extracting data from references cited in systematically identified studies.

